# Prediction of tumor-specific splicing from somatic mutations as a source of neoantigen candidates

**DOI:** 10.1101/2023.06.27.546494

**Authors:** Franziska Lang, Patrick Sorn, Martin Suchan, Alina Henrich, Christian Albrecht, Nina Koehl, Aline Beicht, Pablo Riesgo-Ferreiro, Christoph Holtsträter, Barbara Schrörs, David Weber, Martin Löwer, Ugur Sahin, Jonas Ibn-Salem

**Author notes:** Corresponding authors: Ugur Sahin, and Jonas Ibn-Salem.

## Abstract

Splicing is dysregulated in many tumors and may result in tumor-specific transcripts that can encode neoantigens, which are promising targets for cancer immunotherapy. Detecting tumor-specific splicing is challenging because many non-canonical splice junctions identified in tumor transcriptomes also appear in healthy tissues.

Here, we developed splice2neo to integrate the predicted splice effects from somatic mutations with splice junctions detected in tumor RNA-seq for individual cancer patients. Splice2neo excludes splice junctions from healthy tissue samples, annotates resulting transcript and peptide sequences, and provides targeted re-quantification of supporting RNA-seq reads. We developed a stringent detection rule to predict splice junctions as mutation-derived targets and identified 1.7 target splice junctions per tumor with a false discovery rate below 5% in a melanoma cohort. We confirmed tumor-specificity using independent, healthy tissue samples. Furthermore, using tumor-derived RNA, we confirmed individual exon skipping events experimentally. Most target splice junctions encoded neoepitope candidates with predicted MHC-I or MHC-II binding. Compared to neoepitope candidates derived from non-synonymous point mutations, the splicing-derived MHC-I neoepitope candidates had a lower self-similarity to corresponding wild-type peptides.

In conclusion, we demonstrate that identifying mutation-derived and tumor-specific splice junctions can lead to additional neoantigen candidates to expand the target repertoire for cancer immunotherapies.

**Key Points:**

- splice2neo is a versatile tool for identifying and analyzing of splice junctions as a source of neoantigen candidates
- We identified mutation-retrieved splice junctions supported by RNA-seq in melanoma samples
- The predicted target splice junctions exhibited a strong tumor-specificity as they were absent in healthy tissues.
- Target splice junctions often lead to frame-shift peptides and encode promising neoantigen candidates

## Introduction

Neoantigens are tumor-specific mutated gene products that are presented in form of neoepitopes on major histocompatibility complex (MHC) molecules and that are recognized by CD4+ or CD8+ T cells [1]. Individualized cancer vaccines mediate successful anti-tumor responses by targeting these neoantigens [2–5]. Previous studies mainly focused on targeting neoantigens derived from single nucleotide variants (SNVs) as the most abundant mutation type. However, neoantigens can also derive from other mutation types such as short insertions and deletions (INDELs) or gene fusions which broaden the targetable neoantigen repertoire [6, 7].

During mRNA splicing, multiple isoforms per gene may be created by removal of introns and joining of exons, increasing the functional diversity of the proteome [10]. Importantly, 60% of the alternative splicing events are variable between tissues [11], and inter-individual variability contributes to individual phenotypes [12]. The highly diverse nature of splicing was recently demonstrated by long-read sequencing of more than 70,000 novel transcripts in healthy tissues [13].

Alternative splicing is specifically dysregulated in many tumors, impacting protein function and contributing to tumor heterogeneity [14–17]. For instance, disrupting mutations or expression changes of genes encoding RNA splicing regulators such as SF3B1 alter splicing specifically in tumors [15–19]. Previous studies have shown that indeed novel peptides derived by alternative splicing in tumors can be presented by MHC-I and elicit cytotoxic CD8^+^ T-cell responses in HLA-A24 transgenic mice [18], uveal melanoma [19], acute myeloid leukemia [20] and lung cancer [21]. Alternative splicing can result in frameshifts of the translational reading frame, leading to mutated peptide sequences highly dissimilar to the wild-type proteome. Such dissimilarity could increase the likelihood of T-cell recognition and make them excellent targets.

However, it is challenging to ensure the tumor-specificity of such splicing events. A common strategy is to focus on splice junctions that are absent or low expressed in a set of matched or unmatched healthy tissues [20, 22–24]. However, given the great diversity and stochasticity of alternative splicing across healthy tissue, more evidence is required to consider splice junctions as truly tumor-specific candidates for individualized cancer vaccines [25].

Somatic mutations can directly alter splicing by disrupting canonical splicing motifs or creating novel splicing motifs [26–29]. Notably, integrating somatic mutations with splicing events in whole-genome sequenced pan-cancer cohorts suggested that 34% of the intronic mutations near exon–intron boundaries may affect splicing [27]. Multiple computational tools predicting the effect of mutations on splicing and recent deep learning-based methods considerably improved splicing effect prediction [30, 31].

In contrast to splice junctions detected from RNA-seq alone, expressed splice junctions that are caused by somatic mutations can be considered as truly tumor-specific splice junctions. However, it is currently unclear whether such tumor-specific splice junctions can be predicted reliably from somatic mutations for individual cancer patients and how many splicing-derived neoantigen candidates qualify as promising targets for individualized cancer vaccines.

In this study, we integrated the potential effect of whole exome sequencing (WES) derived somatic mutations on splicing with splicing events detected from RNA-seq, and we identified a small but relevant set of tumor-specific splice junctions that can be interesting targets for individualized cancer vaccines.

## Results

### Splice2neo allows unified analysis of splice junctions from mutation effects and RNA-seq

We developed the R-package splice2neo to facilitate the integration of the effect of somatic mutations on splicing and splicing events detected in RNA-seq for the prediction of splicing-derived neoantigen candidates (Fig. 1A). Splice2neo converts the output of multiple mutation effect prediction tools (MMSplice [32], SpliceAI [33]) and splicing detection tools (SplAdder [34], LeafCutter [35]) in a unified splice junction format, defined by the genomic coordinates of the resulting splice junction positions. Splice2neo is compatible with other R packages and can annotate splice junctions with the resulting transcript sequence, junction position in the transcript sequence, as well as, the resulting peptide sequence. The R-package is designed as a modular library and contains a range of functionalities for customized junction analysis on a joined dataset of multiple samples or for individual samples.

**Fig. 1:**
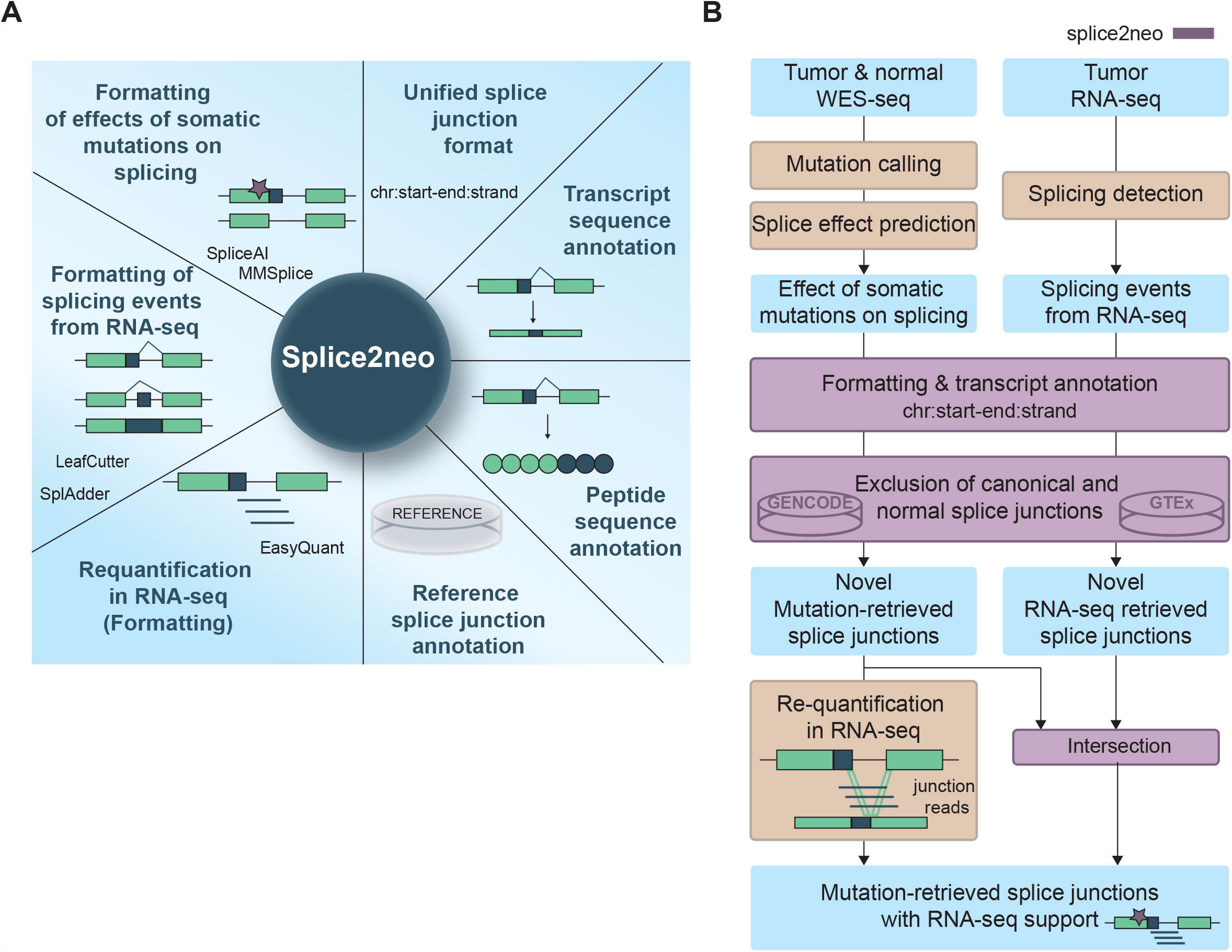
splice2neo is a versatile tool for identifying and analyzing of splice junctions as a source of neoantigen candidates. **(A)** Overview of functionalities implemented in the splice2neo R package. Splice2neo formats the output of several splicing tools into a unified junction format “chr:start-end:strand”, can exclude canonical or normal splice junctions (e.g. from GENCODE or GTEx) and annotates altered transcript and peptide sequences. **(B)** Workflow to detect candidate splice junctions. The effect of somatic mutations on splicing was predicted with SpliceAI and MMSplice, and expressed splicing events were detected with LeafCutter and SplAdder for a given tumor sample, followed by formatting with splice2eno into the unified splice junction format. Novel mutation and RNA-seq retrieved splice junctions were intersected to identify mutation-retrieved splice junctions with RNA-seq support as candidates. To expand the number of candidates, novel mutation-derived were re-quantified in tumor RNA-seq with EasyQuant in a targeted manner.

### Identification of mutation-retrieved splice junctions with support in matched tumor RNA-seq

Somatic SNVs and INDELs are truly tumor-specific as they are detected by comparing WES in tumor and matched normal samples. Such mutations can lead to a loss or gain of splicing donor or acceptor sequence motifs. We reasoned that splice junctions can be defined as tumor-specific targets (1) if they are caused by a somatic mutation, (2) if they are non-canonical and not detected in a large cohort of RNA-seq samples of unmatched healthy tissues [33], and (3) if they are detected in the RNA-seq of the patient’s tumor sample. To develop sufficiently stringent detection criteria for such targets, we analyzed matched tumor and normal WES data and tumor RNA-seq data of 85 melanoma samples from two studies (VanAllen2015 [36], Riaz2017 [37]) with splice2neo as a discovery dataset (Fig. 1B, Fig. 2A). First, we identified between 93 and 10,931 (median 963) somatic SNVs or INDELs in exonic and intronic regions per sample (Fig. 2A, upper panel). Then, we retrieved the potential effects of the identified somatic mutations on splicing with the deep learning-based tools MMSplice [32] and SpliceAI [33]. MMSplice was used to predict if a somatic mutations could cause exon skipping (ES) events and SpliceAI was used to predict all potential 3’ (A3SS) or 5’ (A5SS) alternative splice sites, ES events and intron retention (IR) events. Both tools report effect scores, which reflect the probability or the effect strength of a mutation to alter splicing. Initially, we did not apply cut-offs on these effect scores, and all potential mutation effects on splicing from MMSplice and SpliceAI, including low scoring effects, were annotated with splice2neo and converted into a common splice junction format (“mutation-retrieved splice junction”) (Fig. 2A, middle panel). Next, we removed canonical splice junctions in reference databases (GENCODE) and normal splice junctions previously detected in RNA-seq of 1,740 samples of 53 healthy tissues from the Gene and Tissue Expression (GTEx) atlas [33, 38]. After excluding canonical and normal splice junctions, we considered between 15 and 4,952 (median 430) junctions as novel mutation-retrieved splice junctions for further analysis (Fig. 2A, middle panel).

**Fig. 2:**
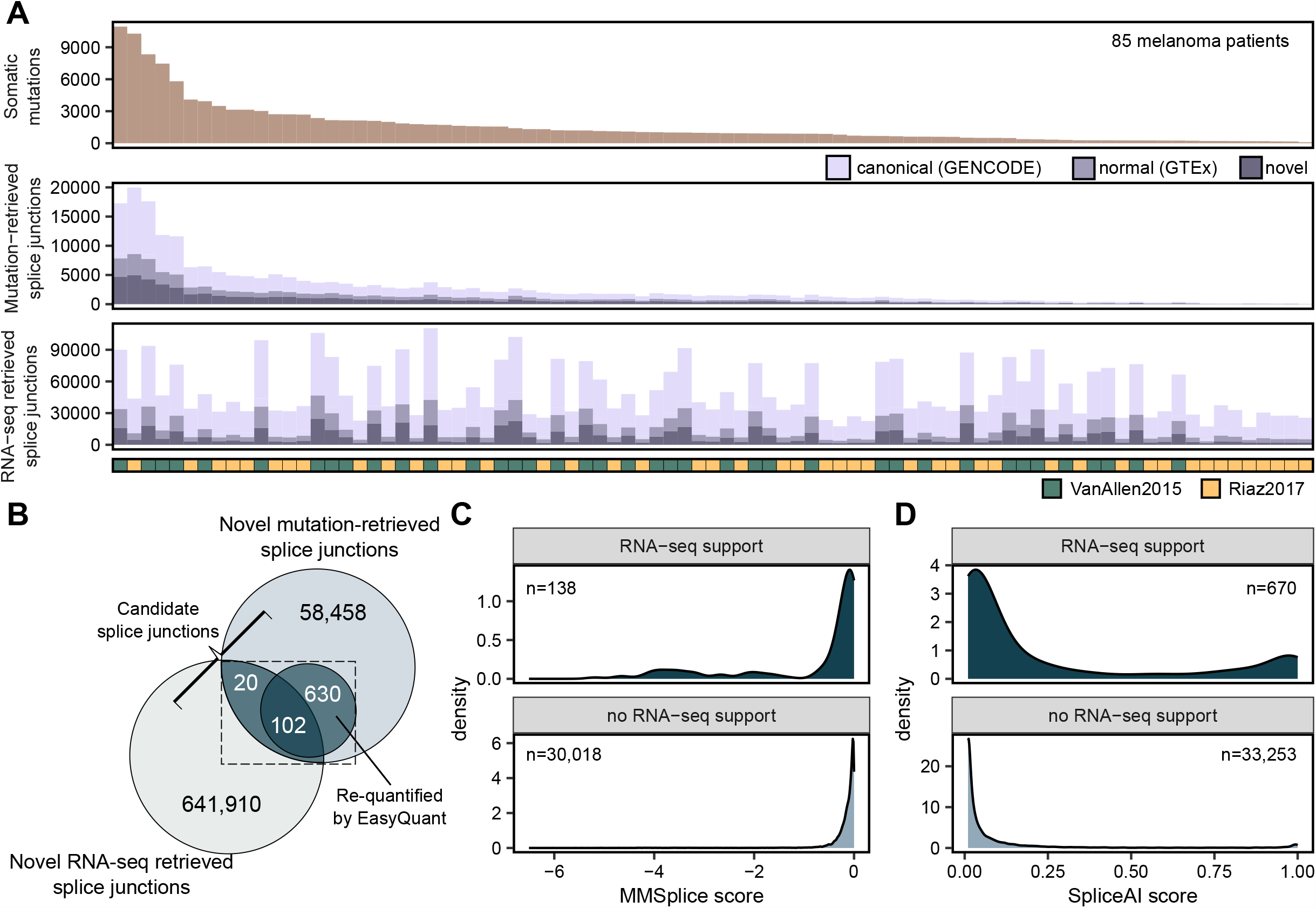
Identification of mutation-retrieved splice junctions supported by RNA-seq in melanoma samples. **(A)** The number of somatic mutations, mutation-retrieved splice junctions and RNA-seq splice derived junctions per sample in the discovery cohort of 85 melanoma samples. **(B)** Overlap of mutation-retrieved splice junctions with those that were found in RNA-seq by SplAdder or LeafCutter and those that were re-quantified in RNA-seq with EasyQuant. Mutation-retrieved splice junctions with RNA-seq support (i.e. by SplAdder, LeafCutter or EasyQuant) were defined as candidate splice junctions. **(C-D)** The distribution of **(C)** MMSplice and **(D)** SpliceAI scores of candidate splice junctions with RNA-seq support and mutation-retrieved splice junctions without RNA-seq support.

The reported effect scores from MMSplice and SpliceAI were low for most of the novel mutation-retrieved splice junctions (Additional file 1: Fig. S1A-B) and decreased with increasing distance of the mutation to the junction position (Additional file 1: Fig. S1C). Although SpliceAI and MMSplice effect scores were correlated for jointly identified ES splice junctions (Pearson correlation R=-0.64), there were many splice junctions with divergent splice effect scores (Additional file 1: Fig. S1D).

Next, we identified splice junctions in matched tumor RNA-seq for the 85 melanoma samples using the RNA-seq-based tools LeafCutter [35] and SplAdder [34]. The number of detected splice junctions differed strongly between the VanAllen2015 and the Riaz2017 cohort (Additional file 1: Fig. S1E-F). Overall, we identified between 16,826 and 110,419 splice junctions per patient, of which between 1,597 and 24,480 were novel (i.e. absent from GENCODE and GTEx) (Fig. 2A, lower panel). The percentage of novel splice junctions was higher among mutation-retrieved splice junctions (26%) than among the splice junctions retrieved from RNA-seq (15%) (Additional file 1: Fig. S1G).

Next, we overlapped the novel mutation-retrieved splice junctions (total n = 59,210) with the novel RNA-seq retrieved splice junctions in RNA-seq (total n = 642,032). For 85 melanoma patients, we identified in total 122 mutation-retrieved splice junctions that were also identified by SplAdder or LeafCutter in RNA-seq (Fig. 2B). We reasoned that targeted re-mapping of RNA-seq reads to pre-defined junction sequences could lead to a higher sensitivity. We developed and implemented the tool EasyQuant and re-evaluated all novel mutation-retrieved splice junctions by quantifying RNA-seq reads supporting the splice junctions or read coverage of retained introns (Fig. 1B, Additional file 1: Fig. S2A). We observed at least one junction read (A3SS, A5SS, and ES) or a median read coverage > 0 (IR) using this targeted approach for 732 out of 59,210 mutation-retrieved splice junctions (Fig. 2B, Additional file 1: Fig. S2B-C). Re-quantified splice junctions and splice junctions identified by SplAdder or LeafCutter had a high overlap of 102 splice junctions (83% of the RNA-seq retrieved splice junctions) (Fig. 2B). Notably, mutation-retrieved junctions supported by RNA-seq were associated with better mutation effect scores from SpliceAI and MMSplice (Fig. 2C-D).

For the identification of target splice junctions that are potentially tumor-specific, we focused on the subset of mutation-retrieved splice junctions with RNA-seq support (n=752) for further analysis.

### Prediction of target splice junctions in melanoma samples

Next, we wanted to identify stringent detection rules to predict which mutation-retrieved splice junctions with RNA-seq support (“candidates”) are caused by a somatic mutation. To estimate the false discovery rate (FDR), we performed a sample permutation analysis by assessing the support of such candidates in RNA-seq data from all other tumor samples without the corresponding somatic mutation. (Fig. 3A, Methods). Here, we observed that half of the candidate splice junctions from ASS or ES events with RNA-seq support in the actual tumor sample were supported by RNA-seq reads in at least one independent tumor sample, resulting in a high estimated FDR of 0.50. This indicates that many of these candidate splice junctions can occur independently of the somatic mutation and, therefore, might not be tumor-specific and that more stringent filtering is required.

**Fig. 3:**
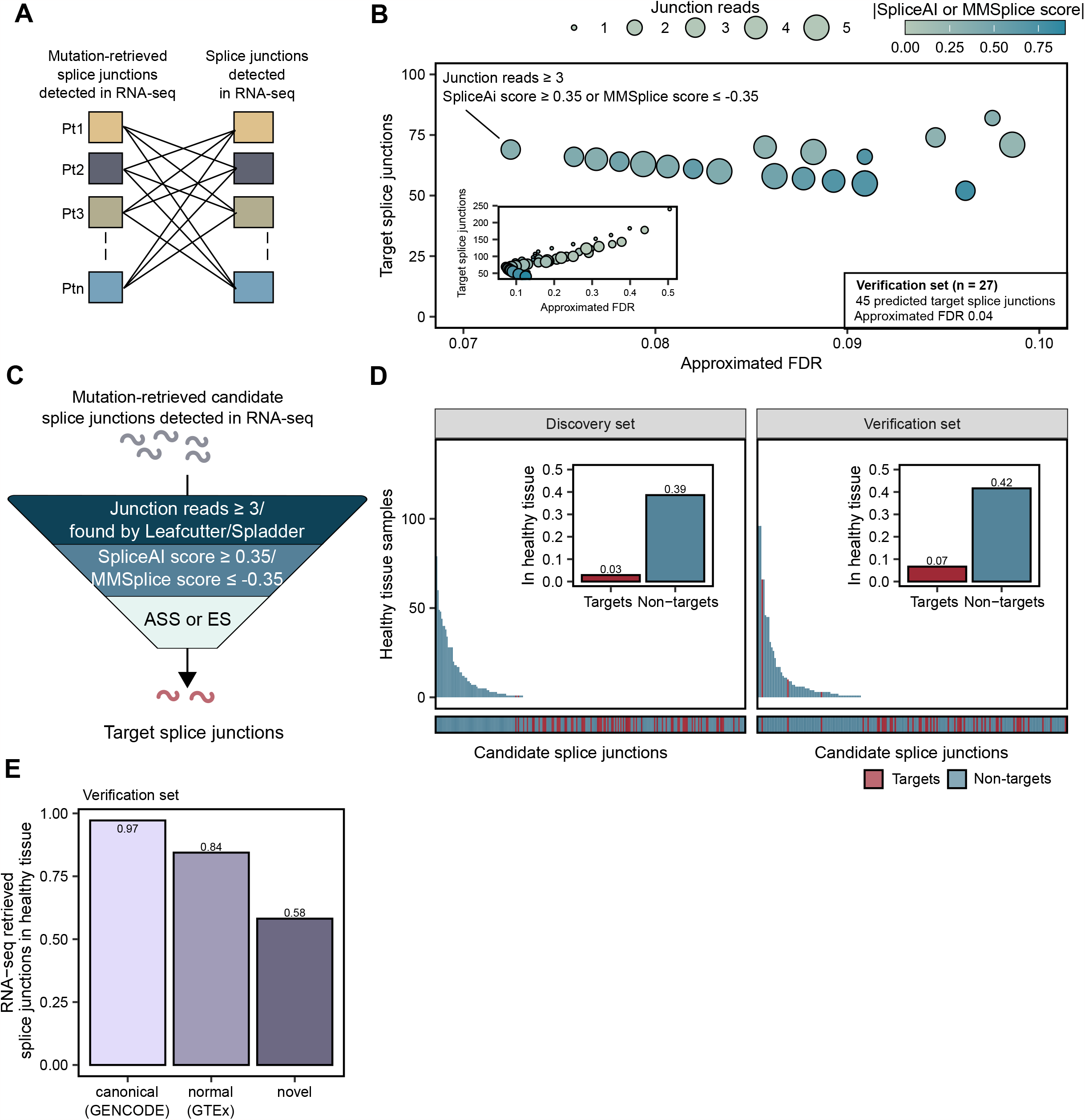
Prediction of tumor-specific target splice junctions based on mutation effect scores and RNA-seq support. **(A)** To calculate the amount of non-specific junctions and estimate a FDR, candidate junctions were compared to splice junctions identified by SplAdder/LeafCutter in other samples’ RNA-seq and their supporting reads were quantified in RNA-seq data of other samples with EasyQuant. **(B)** Candidate splice junctions from the discovery set were gradually filtered by thresholds on the mutation effect scores from MMSplice and SpliceAI, and the re-quantification read support and the resulting estimated FDR and number of target splice junctions was determined. The set of filtering thresholds with lowest estimated FDR was selected as an optimal detection rule for target splice junctions and applied to verification cohort of 27 melanoma samples. **(C)** Target splice junctions were predicted from candidate junctions by the following detection rule: (1) restriction to ES and ASS events, (2) SpliceAI score ≥ 0.35 or MMSplice score ≤ -0.35, and (3) identification by LeafCutter/SplAdder or re-quantification with at least 3 junction reads by EasyQuant. **(D)** Overlap of the target and non-target splice junctions with splice junctions identified by SplAdder/LeafCutter in an additional RNA-seq dataset of 141 samples from 49 healthy tissues in the discovery and verification dataset. **(E)** The fraction of splice junctions derived from RNA-seq found in additional RNA-seq dataset of 141 samples from 49 healthy tissues in the discovery and verification dataset.

Consequently, we gradually filtered candidate splice junctions by thresholds on the mutation effect scores and the re-quantification read support (Fig. 3B). First, we focused on splice junctions from ES and ASS events. We estimated the FDR with the sample permutation analysis and selected the set of filtering thresholds with lowest estimated FDR as optimal detection rule for target splice junctions. This optimal detection rule was defined by (1) SpliceAI score ≥ 0.35 or MMSplice score ≤ -0.35, and (2) identification by LeafCutter or SplAdder or re-quantification with at least 3 junction reads by EasyQuant (Fig. 3B-C). This detection rule resulted in an estimated FDR of 0.07 and identified 69 target splice junctions for 85 melanoma tumors in the discovery cohort (Additional file 2: Table S1). We evaluated the detection rule in an independent verification cohort of melanoma patients (Hugo2016 cohort [39]) and predicted 45 splice junctions for 27 patients with an estimated FDR of 0.04 (Additional file 1: Fig. S3A-B,Additional file 2: Table S1).

For IR events no filter combination led to an estimated FDR lower than 0.50 in the discovery and verification cohort (Additional file 1: Fig. S4), and we, therefore, excluded IR events from further analysis.

To estimate the tumor-specificity of target splice junctions, we compared all candidate splice junctions with splice junctions identified by Leafcutter or Spladder in an additional independent RNA-seq dataset of 141 samples from 49 healthy tissues [7]. Here, only 2 out of 69 (3%) and 3 out of 45 (7%) splice junctions predicted as targets in the discovery and verification set, respectively, were found in any healthy tissue sample. This was a strong reduction compared to splice junctions that were not predicted as targets (39% in discovery set and 40% in verification set) (Fig. 3D). We also used the 141 healthy tissue samples to examine the tumor-specificity of splice junction sets derived from tumor RNA-seq alone. As expected, 97% of the canonical and 84% of the normal RNA-seq derived splice junctions overlapped with the healthy tissue splice junctions. However, also among the novel RNA-seq derived splice junctions, which were already filtered against GTEx, 58% were also found in the independent dataset of healthy tissues (Fig. 3E). Together, this data shows that the here described approach of associating splicing with somatic mutation effects using splice2neo and the detection rule leads to strong enrichment of tumor-specific splice junction targets.

### Experimental confirmation of exon skipping junctions in tumor samples

Next, we experimentally confirmed ES targets by qRT-PCR, which were predicted from eight formalin-fixed, paraffin-embedded (FFPE) tumor samples [7] (Additional file 1: Fig. S5A-B, Additional file 3: Table S2). In total, we tested 21 ES events, of which only one was predicted as a target by the established detection rule (Additional file 1: Fig. S5B-C, Additional file 4: Table S3). This target splice junction (chr20:62606885-62608324:-) was predicted to be caused by a somatic mutation, disrupting an acceptor motif in the gene *SAMD10* (Sterile Alpha Motif Domain Containing 10) (Additional file 1: Fig. S5D). The targeted re-quantification of RNA-seq reads resulted in only two supporting junction reads, but LeafCutter detected the target splice junction in RNA-seq and the associated mutation resulted in the maximal SpliceAI score of 1.0 (Additional file 1: Fig. S5D-E, Additional file 4: Table S3), indicating a strong effect on splicing.

The expression of the target splice junction and two additional non-target junctions could be confirmed with qRT-PCR on RNA level (Additional file 1: Fig. S5E, Additional file 4: Table S3).

### Splice junctions can generate neoantigen candidates

Next, we examined whether the predicted target splice junctions encode novel peptide sequences and therefore qualify as neoantigen candidates. For in-frame junctions, we considered the peptide sequence (-/+13 amino acids) around the junction and potentially inserted amino acids using splice2neo. For frame-shift junctions, the peptide sequence was extended until the next stop codon. We refer to peptide sequences from predicted tumor-specific targets as “neoantigen candidates”. In total, 42 of the 45 predicted target splice junctions affected coding sequences and were translated into at least one peptide sequence, resulting in 45 neoantigen candidates for the 27 melanoma patients (Additional file 5: Table S4). Depending on the mutation load, we identified in average 1.7 neoantigen candidates from alternative splicing per melanoma patient (Fig. 4A). We found that the majority of neoantigen candidates in the verification cohort derived from an A3SS event (42%), followed by ES events (31%) (Fig. 4B). Furthermore, 62% of splicing-derived neoantigen candidates were generated by a frameshift. While also in-frame A3SS or A5SS events could generate neoantigen candidates longer than 26 amino acids if additionally amino acids were inserted, frame-shift splice junctions led generally to longer novel peptide sequences of up to 134 amino acids (Fig. 4C-D).

**Fig. 4:**
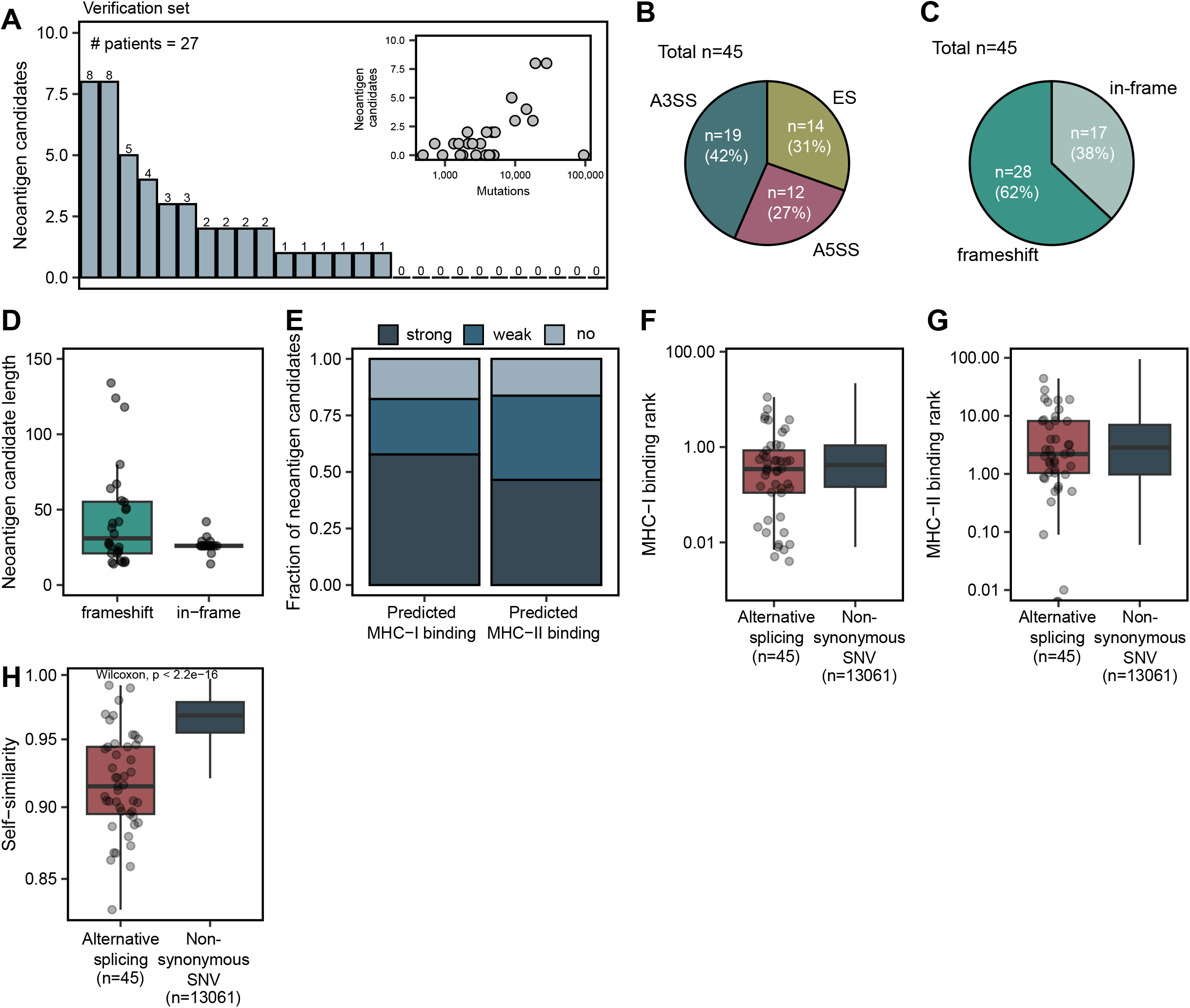
Target splice junctions generate neoantigen candidates. **(A)** The number of neoantigen candidates per sample in the verification dataset. Inlet: Correlation of the neoantigen candidate count with the tumor mutation burden. **(B)** The fraction of neoantigen candidates from A3SS, A5SS and ES events. **(C)** The fraction of frameshift and in-frame neoantigen candidates. **(D)** The length distribution of frameshift and in-frame neoantigen candidates. **(E)** The fraction of strong or weak binding MHC-I and MHC-II neoantigen candidates.(strong: MHC-I binding rank < 0.5, MHC-II binding rank < 2, weak: 0.5 ≤ MHC-I binding rank < 2, 2 ≤ MHC-II binding rank < 10, no: MHC-I binding rank ≥ 2, MHC-II binding rank ≥ 10) **(F-G) (F)** MHC-I and **(G)** MHC-II binding rank of the best predicted neoepitope per neoantigen candidate for those from alternative splicing (AS) and SNVs. SNV-derived neoantigen candidates were retrieved as described in [69] **(H)** Self-similarity [41] of the best predicted neoepitope per neoantigen candidate for those from alternative splicing and those from SNVs.

Next, we predicted MHC-I and MHC-II binding features with NeoFox [40] using patient-specific MHC-I and MHC-II alleles. We found that 58% of the neoantigen candidates from alternative splicing were predicted to generate at least one strongly binding MHC-I epitope (MHC-I binding rank < 0.5), and 47% of the neoantigen candidates were predicted to generate at least one strongly binding epitope for MHC-II (MHC-II binding rank < 2) (Fig. 4E). Neoantigen candidates from alternative splicing had comparable MHC-I and MHC-II binding ranks to neoantigen candidates derived from non-synonymous SNVs of the same cohort (Fig. 4F-G). However, the best-predicted MHC-I neoepitope per neoantigen candidate from alternative splicing was less self-similar to the wild-type proteome [41] compared to the best-predicted neoepitope per neoantigen candidate from nonsynonymous SNVs (Fig. 4H), indicating a stronger potential for immunogenicity in the group of neoantigen candidates derived from alternative splicing.

## Discussion

Disruption of canonical splicing in tumors can generate novel gene products that might be excellent targets for individualized cancer vaccines [6, 42]. While RNA-seq can detect thousands of novel splice junctions in individual tumor samples, it is challenging to ensure their specificity to tumor cells [25, 43]. One approach is to predict neoantigen candidates from alternative splicing by combining RNA-seq from tumor samples with RNA-seq from matched healthy tissue [22]. Given the diversity of splicing across tissues and cell types, it is questionable which healthy tissue is suitable as a control and if a single normal sample is sufficient. Alternatively, tumor-specificity might be defined by the absence of splice junctions from large healthy tissue sample collections of unrelated subjects [23, 24]. Databases of canonical reference splice junctions or large RNA-seq sample collections of no healthy tissues, such as GTEx [38], are valuable resources for constructing exclusion lists. However, it remains unclear if such data is suitable to capture splicing variations in rare cell types or conditions as well as individual splicing events caused, for example, by rare germline mutations.

In this study, we hypothesized that alternative splicing created by a loss or gain of canonical splicing sequence motifs by somatic mutations are eligible as tumor-specific targets and might be suitable neoantigen candidates for individualized cancer immunotherapy approaches. With splice2neo, we implemented an analysis approach to identify mutation-retrieved splice junctions with support in RNA-seq as candidates for potentially tumor-specific targets for individual cancer patients.

Multiple other tools were recently described to predict neoantigen candidates from alternative splicing. NeoSplice [22] and ASNEO [23] rely on a single RNA-seq-based method for splicing detection and are end-to-end pipelines from raw files to neoantigen candidates. IRIS [24] is a modular neoantigen prediction pipeline based on RNA-seq and supports customized pipelines. Regtools [28] provides functionalities to integrate DNA sequencing and RNA-seq data to identify potential splice-associated variants. DICAST [44] integrates several RNA-seq splicing tools for unified junction analysis. Both, Regtools and DICAST lack the transcript and protein sequence annotation for functional downstream analyses. In contrast to static end-to-end pipelines, splice2neo employs third-party tool for splicing prediction and is designed as a modular library of multiple functionalities for customized splice junction analysis. It includes the unified integration of results from upstream tools, exclusion of canonical or normal junctions, and annotation with transcript and peptide sequences. The implementation of splice2neo as an R-package allows compatibility with multiple other methods from the Bioconductor project [45, 46] for interactive analysis or integration into target identification pipelines. Splice2neo is not limited to the currently supported upstream tools but can be easily extended in the future to support other tools for alternative splicing detection or other event types, such as splicing junctions between exons and transposable elements [21, 47]. The proposed approach works with data from single patients, which is relevant in the context of neoantigen prediction for individualized cancer vaccines but might also be used to analyze larger cohorts of tumor samples.

We initially considered all novel mutation-retrieved splice junctions that were supported by RNA-seq without any cut-off on splice effect scores. Only if we applied a stringent detection rule on the mutation effect scores and RNA-seq support, we were able to decrease the FDR and enrich for expressed splice junctions that could indeed be associated with somatic mutations. Depending on the somatic mutation burden, on average 1.7 neoantigen candidates from alternative splicing could be predicted per patient with an estimated FDR of 0.04. We assume that the number of neoantigen candidates from somatic mutation-derived splicing is not many magnitudes higher but still can be increased by technical advances. In the future, the computational analysis of larger tumor cohorts from more diverse entities and using personalized reference genomes may allow more sensitive detection rules, e.g., by a machine learning approach.

Studies predicting splicing-derived neoantigen candidates from tumor RNA-seq alone or in combination with matched normal RNA-seq reported markedly higher numbers of non-canonical splice junctions per patient in several tumor entities [17, 21, 22]. However, it remains unclear whether those splice junctions identified from RNA-seq alone are truly tumor-specific and qualify as safe and effective targets in individualized cancer vaccines.

We were not able to show that candidate splice junctions from IR events are potentially tumor-specific. IR detection can be generally challenging as RNA-seq reads might derive from unspliced pre-mRNA, repeat regions, or other transcripts [48, 49]. However, it was shown that somatic mutations indeed can cause IR events [29], that IRs can generate MHC-I binding epitopes [50] and that the load of non-canonical peptides derived from IRs correlated with favorable prognosis in pancreatic cancer [51]. These observations suggest that also IRs could contribute to the neoantigen repertoire. Improved computational tools for IR detection and long read sequencing might allow to better investigate splice junctions from IRs in the future [18, 48, 49, 52].

Here, we showed that the predicted target splice junctions from somatic mutations can generate neoantigen candidates. By association with somatic mutations, the number of these neoantigen candidates depends on the tumor mutation burden. Therefore, neoantigen candidates from mutation-derived splicing might be in particularly rare in tumor entities with low tumor mutation burden which still could be sufficient for anti-tumor response as long as they are of high quality as shown in pancreatic cancer [53, 54]. Indeed, those targets frequently cause frameshifts and lead to longer novel peptide sequences, leading to strong MHC binding neoepitope candidates that are dissimilar to the wild-type proteome. These features may characterize splicing-derived neoantigen candidates as promising targets for individualized cancer vaccines [1]. Future studies are required to examine if the identified neoantigen candidates indeed mount functional T-cell responses upon individualized cancer vaccines.

## Methods

### Whole exome sequencing and somatic mutation calling

Whole exome sequencing and somatic mutation calling is described in the Supplementary Methods.

### Predicting the effect of somatic mutations on splicing

The effect of somatic mutations on splicing was predicted with SpliceAI [33] (v1.3.1) and MMSplice [32] (v2.1.1). For SpliceAI, the GENCODE V24 canonical annotation (grch37) included in the package was used. GENCODE v34lift37 was used as annotation for MMSplice. Both tools were run with default settings.

### Identifying alternative splicing events from RNA-seq

RNA-seq reads were aligned to the hg19 reference genome with STAR [63] (v2.7.0a). The options ‘twopassMode Basic --outSAMstrandField intronMotif’ were used to generate BAM files for the tool LeafCutter based on recommendation in the manual. Alternative splicing events were detected with SplAdder [34] (v3.0.2) and LeafCutter [35] (v0.2.9). Splicing events from exon skipping, alternative 3’ or 5’, intron retention and mutually exclusive exons were identified with SplAdder, ignoring mismatch information in the alignment file and using GENCODE (v34lift37) annotation. Splicing events were identified with LeafCutter annotation-free using the settings as recommended in the manual (-m 50 -l 500000 --checkchrom). Here, the tool how_are_we_stranded_here [64] (v1.0.1) was used to computationally determine the strandedness of the RNA-seq and regtools [28] (v0.5.2) was used to extract junctions using the recommended parameters of -a 8 -m 50 -M 500000. The union of RNA-seq derived junctions was considered per sample if RNA-seq replicates were available.

### Splice2neo: prediction of splice junctions and derived mutated transcript and peptide sequences

We developed the R-package splice2neo (https://github.com/TRON-Bioinformatics/splice2neo) to identify mutation-retrieved splice junctions with RNA-seq support defined (see Supplementary methods for a detailed description). We used splice2neo to convert the raw output from LeafCutter, SplAdder, MMSplice and SpliceAI into resulting splice junctions. While converting the output of SpliceAI with splice2neo, only transcripts related to the annotated gene were considered. Using splice2neo, we further annotated splice junctions with the resulting modified transcript and peptide sequences. Furthermore, splice junctions were annotated whether they are canonical using a database of canonical splice junctions built from GENCODE v34lift37 using the R-package GenomicFeatures [65] (v.1.46.5) and classified as “normal” if they were contained in a dataset of normal splice junctions previously identified by Jaganathan et al. with LeafCutter in 1,740 RNA-seq samples of 53 healthy tissues from GTEx [33]. We used splice2neo v0.6.2 in this study.

### Re-quantification of RNA-seq reads for mutation-retrieved splice junctions

To quantify the number of supporting RNA-seq reads for given splice junctions in a sensitive and targeted manner, we developed and applied EasyQuant (v0.4.0, https://github.com/TRON-Bioinformatics/easyquant, see Supplementary methods for a detailed description). EasyQuant was run in the interval mode (--interval_mode) and reads covering the junction position by at least 10bp were considered to support the splice junctions (-d 10). RNA-seq reads were mapped to the context sequence with STAR [63] (-m star). The following STAR parameters were provided in the config file of Easyquant in the general section mismatch_ratio=0.015 (outFilterMismatchNoverReadLmax), indel_open_penalty=-1000 (scoreDelOpen), indel_extension_penalty=-2 (scoreInsOpen). If RNA-seq replicates were available, read counts and summary values were summed up per splice junction. To not overestimate intron retention read support, we considered here only potential intron retention events for which the retained intron region did not overlap with any exon of any other transcript.

### False discovery rate estimation for prediction of target splice junctions

We aimed to derive a detection rule to predict splice junctions derived from somatic mutations as tumor-specific targets in a discovery cohort of 85 melanoma patients (VanAllen2015 [36], Riaz2017 [37]) and to confirm it in a verification cohort of 27 melanoma patients (Hugo2016 [39]).

To identify potential false positive splice junctions, we compared the RNA-seq support of novel mutation-retrieved splice junctions from the actual sample with the RNA-seq support from a different individual that does not has the corresponding mutation. The permutation analysis was performed among all samples per study (e.g. within VanAllen2015 and within Riaz2017 separately). Predicted splice junctions from somatic mutations were re-quantified with EasyQuant in the RNA-seq of all other independent tumor samples. Second, mutation-retrieved splice junctions were searched among the RNA-seq derived junctions identified in other biological independent samples.

A mutation-retrieved splice junction with RNA-seq support was labelled as false positive if there was support by re-quantification or SplAdder/LeafCutter derived splice junctions in at least one other independent sample. The estimated FDR was defined as the number of such false positives divided by all considered candidate splice junctions in the discovery or verification cohort.

To identify an appropriate detection rule, candidate splice junctions were gradually filtered by a grid of thresholds on the mutation effect scores from MMSplice and SpliceAI [0, 0.05, 0.1, 0.15, 0.2, 0.25, 0.3, 0.35, 0.4, 0.5, 0.6, 0.7, 0.8, 0.9], and the re-quantification read support [1, 2, 3, 4]. The number of remaining splice junctions and the FDR was estimated for each threshold combination. The threshold combination with the lowest FDR in the discovery cohort was defined as the detection rule and remaining candidate splice junctions were defined as targets. This detection rule was applied to the verification cohort to estimate the resulting number of target splice junctions and the final estimated FDR.

### Annotation of neoantigen candidates

MHC -I and –II genotypes were determined with HLA-HD [67] (v1.2.0.1) for each patient based on the WES-seq data of the normal sample. NeoFox [40] (v1.0.2) was used to annotate neoantigen candidates derived by target splice junctions with neoantigen features. Here, the predicted mutated peptide sequences without information about variant allele frequency or transcript expression was provided as input to NeoFox. Furthermore, MHC-I and -II genotypes of the patients were provided to the tool as input.

### Confirmation of splice junction expression with qRT-PCR

Primer design and confirmation of splice junctions with qRT-PCR is described in Supplementary Methods.

## Declarations

### Ethics approval and consent to participate

The patient material of the FFPE cohort was collected as part of the RB_T002 research program (DRKS-ID: DRKS00011790). The studies were carried out in accordance with the Declaration of Helsinki and good clinical practice guidelines and with approval by the institutional review board or independent ethics committee of each participating site and the competent regulatory authorities. All patients provided written informed consent.

### Availability of data and materials

RNA-seq data for the FFPE cohort [7] is available in European Genome-phenome Archive (EGA) under accession number EGAS00001004877 and WES was uploaded as EGAS00001007589. Data for the Hugo cohort [39] is available in the Sequence Read Archive (SRA) under accession numbers SRP067938, SRP090294 (WES-seq) and SRP070710 (RNA-seq). Data for the Riaz cohort [37] is available under accession numbers SRP095809 (WES-seq) and SRP094781 (RNA-seq). Data for the Van Allen cohort [36] is available in dbGap under accession number phs000452.v2.p1. RNA-seq for the cohort of healthy tissue is available under is in SRA under NCBI BioProject ID PRJNA764684.

The source code and documentation of splice2neo (https://github.com/TRON-Bioinformatics/splice2neo) and EasyQuant (https://github.com/TRON-Bioinformatics/easyquant) are available under open source licenses on GitHub.

### Competing interests

U.S. is co-founder, chief executive officer and stock owner of BioNTech SE. The remaining authors declare no competing interests.

### Funding

This work was supported by an ERC Advanced Grant to U.S. (ERC-AdG 789256).

## Authors’ contributions

Conceptualization: JI. Methodology: FL, JI. Software: FL, PS, PRF, JI. Data curation: FL, PS. Formal analysis: FL, JI, Investigation: FL, PS, MS, AH, CA, NK, AB, PRF, CH, DW, JI. Supervision: BS, DW, ML, US, JI. Visualization: FL, JI. Writing—original draft: FL, JI. Writing—reviewing and editing: FL, BS, DW, ML, US, JI. All authors read and approved the final manuscript.

## Acknowledgements

We thank the RB_T002 research program (DRKS-ID: DRKS00011790) patients, from whom analyzed samples were obtained, and we thank the involved study site teams for their support and collaboration. We thank O. Akilli-Oeztuerk for support with biosampling. This work was supported by an ERC Advanced Grant to U.S. (ERC-AdG 789256). The authors further acknowledge the authors and generators of datasets used in this work and the grants that supported the studies.

## Notes

### Summary of Updates

The manuscript text has been improved and shortened for calrity. Figure S5A has been updated to fix a minor error. Affiliations have been updated.

