## Additional file 1 for "Prediction of tumor-specific splicing from somatic mutations as a source of neoantigen candidates"

### Supplementary Methods

#### Whole exome sequencing

Genomic DNA was isolated from 10µm curls of FFPE tissue using the QIAamp DNA FFPE Tissue Kit (Qiagen) or Maxwell RSC DNA FFPE Kit (Promega) according to the manufacture's protocol. The DNA quantity was assessed using Qubit 3 fluorometer using the dsDNA HS Assay Kit (Invitrogen).

Whole exome libraries were prepared in duplicates with an input of 100 ng genomic DNA from FFPE tissue samples. The genomic DNA was fragmented to a size range of 200 ± 50 bp using the Covaris S220 instrument. The fragmented DNA was end-repaired and adenylated followed by ligation of sequencing adaptor and the appropriate eight-nucleotide NEXTFLEX DNA barcode (Perkin Elmer) and pre-amplification of the library using the KAPA Hyper Prep kit (Roche). Subsequently, target regions were hybridized to biotinylated baits (SureSelectXT Human All Exon v6 and SureSelect XT Reagent kit, Agilent) and isolated using streptavidin-coated magnetic beads (Invitrogen). The post-capture library was amplified with the KAPA Library Amplification kit (Roche). Final library quantity and quality was assessed using the Qubit 3 fluorometer with Qubit dsDNA HS Assay Kit (Invitrogen) and the BioAnalyzer with High Sensitivity DNA Kit (Agilent).
All libraries were sequenced in paired-end mode (2 x 50 nt) on an Illumina NovaSeq 6000 instrument resulting in around 150 million distinct sequencing reads per library.

#### Somatic mutation calling

For WES read alignment and somatic mutation calling, we used the tronflow pipelines [55–57] implemented in Nextflow [58]. In short, reads from WES tumor and normal sequencing were aligned to the reference genome hg19 using bwa [59] (v0.7.17). Base score recalibration was performed with GATK4 [60] (v4.2.0.0) and duplicated reads were removed with Picard (GATK4 v.2.0.0). Single nucleotide variants (SNVs) and short insertions and deletions (INDELs) were detected with Mutect2 [61] (GATK4 v.2.0.0) without restriction to any target region and using the GnomAD [62] as a resource of germline mutations. Only mutation calls with PASS in the filter column were considered and calls from individual replicates were combined per sample.

#### Splice2neo: prediction of splice junctions and derived mutated transcript and peptide sequences

We developed the R-package splice2neo (https://github.com/TRON-Bioinformatics/splice2neo) to identify splice junctions defined by the chromosome (chr), the last genomic position of the left exon (start), the first base of the right exon (end) and the transcriptional direction (strand) in the format "chr:start-end:strand" directly from the output of multiple tools. The R-package relies mainly on the functionalities of the Bioconductor packages GenomicRanges [65] and Biostrings [66]. Splice2neo converts the raw output from LeafCutter, SplAdder, MMSplice and SpliceAI into resulting splice junctions.

LeafCutter and SplAdder provide the genomic coordinates of the identified splicing events. Splice2neo transforms all events into the standardized junction format covering all potential splice junctions including canonical junctions.

SpliceAI predicts the probability of a mutation on splicing and provides the position of the alternative splicing events relative to the mutation. Table 1 describes the rules on how splice2neo creates splice junctions based on the predicted effect by SpliceAI. Mutations can be covered by more than one gene and the splice effect of a somatic mutation may differ between genes depending of the position of the somatic mutation within the respective gene. Splice2neo optionally considers only transcripts related to the gene annotated by SpliceAI while predicting the effect on splicing.

MMSplice predicts the change on percent spliced in (PSI) for a given annotated exon. Splice2neo considers events with PSI score <= 0 as exon skipping events and uses the delta_logit_psi output as mutation effect score. Here, splice junctions are built by the end of the upstream exon and the start of the downstream exon.

**Table 1: Rules on how splice junctions are created based on the mutation effect predicted by SpliceAI.**

| Change | Class | Left junction coordinate | Right junction coordinate |
|---|---|---|---|
| **Donor loss** | Intron retention | pos | pos + strand_offset |
| **Donor loss** | ES | Upstream_end | downstream_start |
| **Donor gain** | A5SS | pos | downstream_start |
| **Acceptor loss** | Intron retention | pos – strand_offset | pos |
| **Acceptor loss** | ES | Upstream_end | downstream_start |
| **Acceptor gain** | A3SS | Upstream_end | pos |

The table describes the rules on how the left and right coordinate of the splice junction ("chr:start-end:strand") are determined based on the predicted effect from SpliceAI. Pos refers to the position that is predicted to be affected by the mutation by SpliceAI. Upstream_end and downstream_start refer to exon end or start coordinates. Splice junctions that relate to an intron retention event are defined by the rule `chr:pos-(pos+1):strand` and must cover an exon-intron boundary. The strand_offset is +1 for transcript on the positive strand and -1 otherwise.

Splice2neo, provides function to annotate splice junctions with the resulting modified transcript and peptide sequences.

The transcript context sequence covers by default 200-bp exonic sequence up- and downstream of a splice junction from an A3SS, A5SS or ES event ("cts_seq"). In case of intron retention events, the sequences covers the complete intronic sequence, flanked by 200-bp (default) exonic sequence up-and downstream of the intron retention event. The position of the splice junction in the sequence or the intron interval are given by "cts_junc_pos". A unique id ("cts_id") is given based on the context sequence and position as a XXH128 hash value.

Given the splice junction and coordinates of coding sequences (CDS) of reference transcripts, splice2neo annotates the resulting protein sequence for each affected transcript and the relevant peptide context sequence. In cases of in-frame events, this peptide context sequence covers junctions with potentially novel amino acid sequences (residue at the junction position or inserted novel residues) flanked with 13 wild-type amino acids up- and downstream. Frameshift peptides are flanked by an upstream wild-type region and are translated until the next stop codon. If splice junctions do not generate mutated gene products, no peptide context sequence is provided. This can be the case if junction does not affect the CDS, or the "mutated" gene product is only a truncated version of the wild-type CDS.

We used splice2neo v0.6.2 in this study.

#### Re-quantification of RNA-seq reads for mutation-retrieved splice junctions

To quantify the number of supporting RNA-seq reads for given splice junctions in a sensitive and targeted manner, we developed and applied EasyQuant (v0.4.0) https://github.com/TRON-Bioinformatics/easyquant. Analogous to the re-quantification for gene fusions that we previously developed in EasyFuse [7], EasyQuant aligns reads to a context sequence ("cts_seq") constructed from the splice junction and calculates the reads supporting the junction (Supplemental Fig. 1). RNA-seq reads are mapped to the context sequence with STAR [63] with the following parameters: --outFilterMismatchNoverReadLmax 0.015 --alignEndsType EndToEnd --outFilterMultimapNmax -1 --outSAMattributes NH HI AS nM NM MD --scoreDelBase -2 –scoreInsBase -2. The STAR parameter scoreDelOpen, scoreInsOpen and outFilterMismatchNoverReadLmax can be defined by the user in the configuration file of Easyqunt by providing indel_open_penalty, indel_extension_penalty and mismatch_ratio, respectively in the section “general”. Next, the context sequence is divided into intervals based on the provided positions. For each interval, the number of reads within the interval is counted as well as reads overlapping interval end positions. Junctions from alternative splice sites or exon-skipping events are divided into two intervals. The end of the first interval represents the splice junction of interest. Reads covering the junction position by at least 10bp are considered to support the splice junction (“junction reads”). Context sequences for intron retentions are divided into three intervals, whereby the second interval is the retained intron. We summarized the coverage of the entire intron by the median number of reads per position in the interval. The output values from EasyQuant are defined in Table 2.

**Table 2: Read quantification in EasyQuant per interval and input sequence.**

| Value | Definition |
|---|---|
| **overlap_interval_end_reads** | Number of reads that overlap interval end position by at least 10bp |
| **span_interval_end_pairs** | Number of read pair that span the interval end. Each read of the pair must be map on a different interval. |
| **within_interval** | Number of read that maps completely within the interval. |
| **coverage_perc** | Percent of interval positions that are covered by at least one read read. |
| **coverage_mean** | Mean number of mapped reads per interval position. |
| **coverage_median** | Median number of mapped reads per interval position. |

If RNA-seq replicates were available, read counts and summary values were summed up per splice junction. To not overestimate intron retention read support, we considered here only potential intron retention events for which the retained intron region did not overlap with any exon of any other transcript.

#### Confirmation of splice junction expression with qRT-PCR

Primers were designed as described previously [7]. In short, context sequences as predicted by splice2neo were used for primer design with Primer-BLAST [68]. Primer design was guided by the following restrictions: (1) no primer should align within 20 bp upstream and downstream of the junction; (2) amplicons must span the junction; and (3) amplicon size should not exceed 150 bp. Primer pairs aligning to off-target loci were removed.

Splice junctions were validated in previously extracted total RNA from fresh frozen tissue samples [7]. The RNA was reverse transcribed into cDNA using TAKARA PrimerScript RT Reagent Kit with gDNA Eraser. Reverse Transcriptase was replaced with water in no amplification controls.

qRT-PCR was performed on a BioRad CFX384 instrument using BioRad SsoAdvanced Universal SYBR Green Supermix. After polymerase activation for 30 s at 98°C, qPCR were run in 40 cycles of two-step PCR with denaturation for 10 s at 98°C and annealing/elongation for 30 s at 60°C.

The PCR products were analyzed for the presence of amplicons matching the expected sizes using QIAGEN's QIAxcel Advanced capillary gel electrophoresis instrument with a 15 bp/600 bp alignment marker.

A custom R script was used to evaluate qRT-PCR and capillary gel electrophoresis results and to assign each tested amplicon as positive or negative given a defined set of rules. (1) The amplicon had to be detected in qRT-PCR with a Ct value below or equal to 35 and at least 5 cycles earlier compared to water and no amplification controls. (2) The amplicon product size had to match the expected amplicon size (allowing a maximum difference of 15% in amplicon size). (3) The amplicon associated peak had to cover at minimum 15% height relative to the total peak heights. Amplicons fulfilling the first two rules but with a relative height below 15% were classified manually by an expert user.

#### Read support visualization with sashimi plots

A Sashimi plot (Additional File: Fig. S5D) was generated with the python package rmats2sashimiplot (v 2.0.4) (https://github.com/Xinglab/rmats2sashimiplot) using RNA-seq BAM files with coordinate and GENCODE (v34lift37) annotation. The original Sashimi plot was manually modified to include one further affected transcript while a non-affected transcript was removed from the plot.

### Supplementary Figures

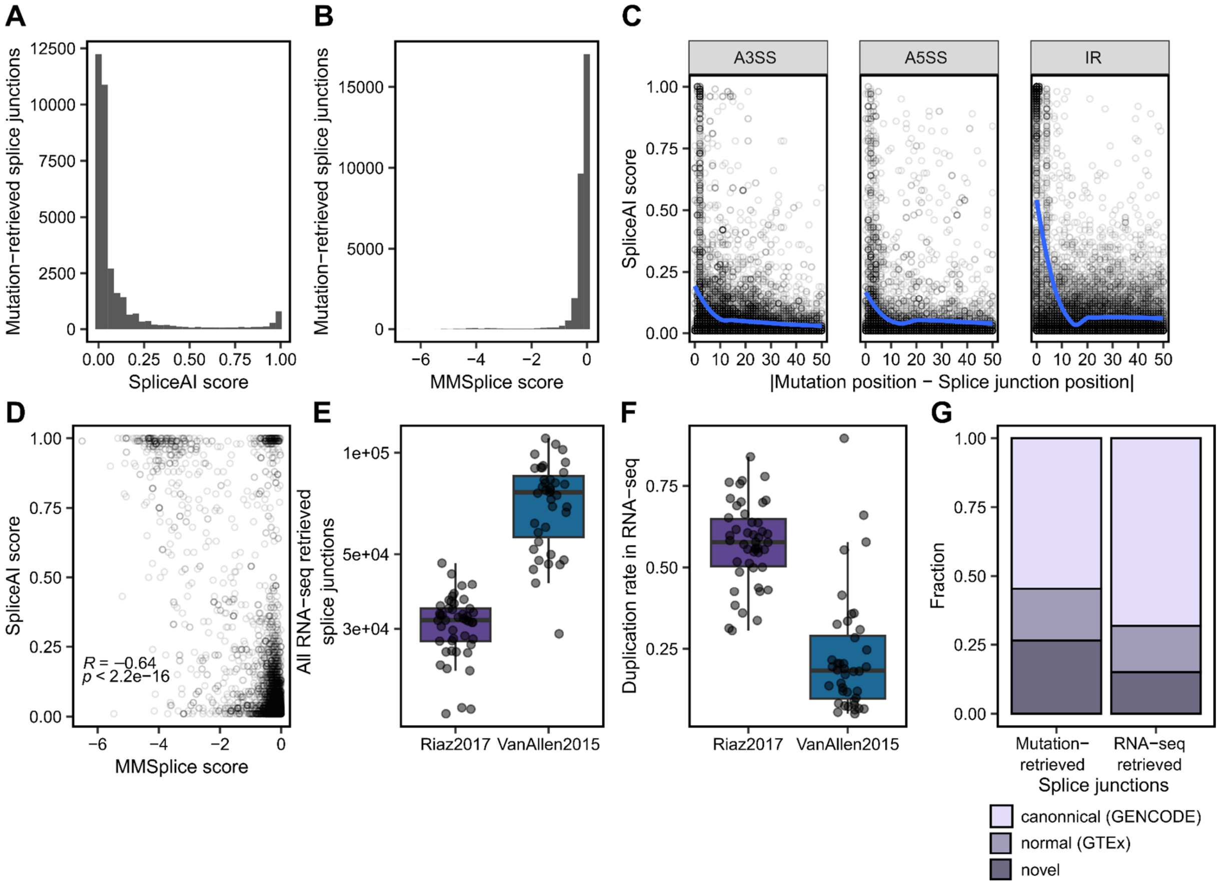


**Fig. S1: Overview of the discovery set. (A-B)** The distribution of **(A)** SpliceAI and **(B)** MMSplice scores in the discovery set. **(C)** The distribution of SpliceAI score dependent on the genomic distance between splice junctions from A3SS, A5SS and IR events and the mutation they were retrieved from. **(D)** Correlation of MMSplice and SpliceAI score for splice junctions from exon skipping events. Negative MMSplice scores indicate stronger exon exclusion effects. **(E)** The number of non-canonical junctions derived from RNA-seq per sample in the Riaz2017 and VanAllen2015 cohorts. **(F)** The duplication rate in RNA-seq per sample in the Riaz2017 and VanAllen2015 cohorts. **(G)** The fraction of splice junctions found in GENCODE and GTEx, found in GENCODE or GTEx and novel junctions among splice junctions derived from RNA-seq and mutation-derived splice junctions.

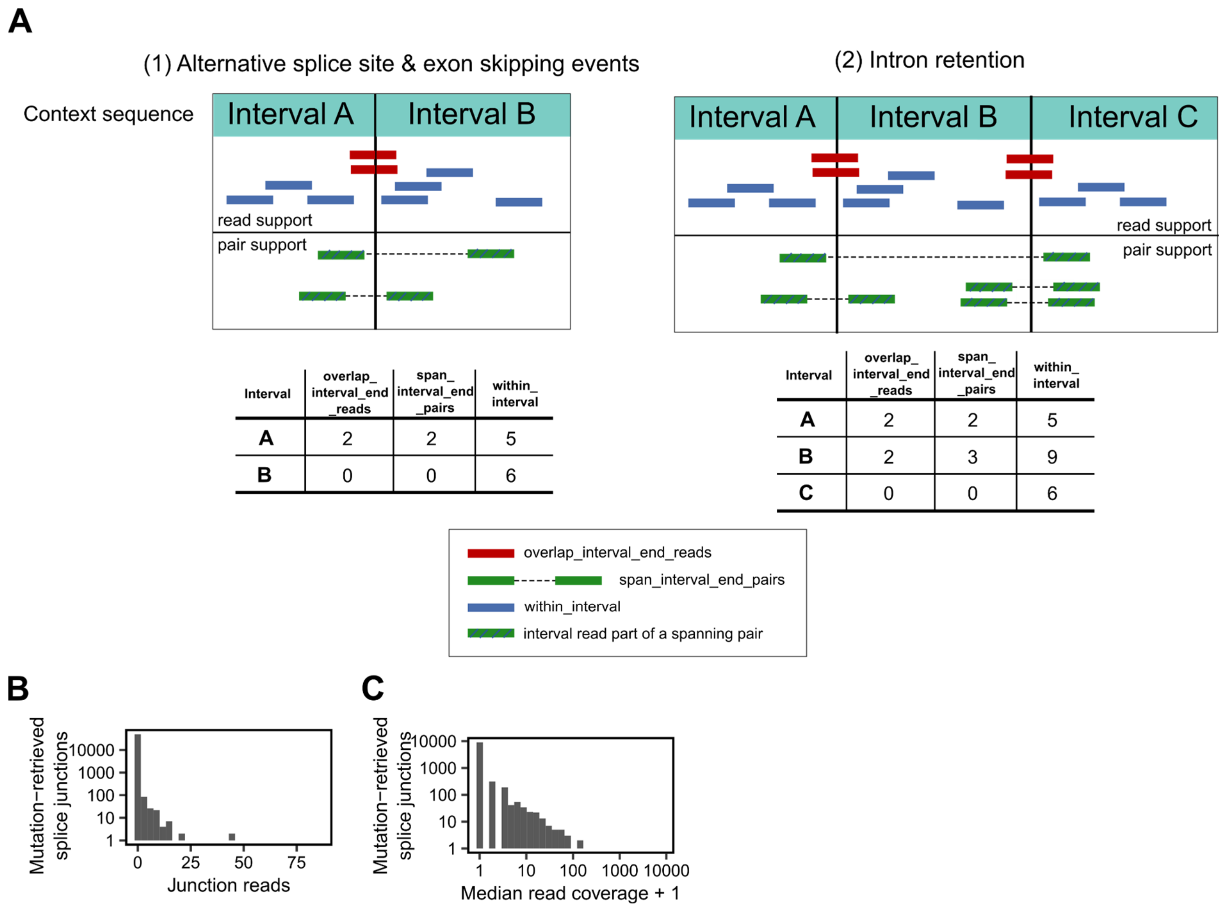


**Fig. S2: Re-quantification with EasyQuant in the interval mode. (A)** EasyQuant divides a given context sequence into intervals based on provided coordinates. Both context sequence and interval coordinates can be determined with splice2neo. Upon read alignment to the context sequence, EasyQuant counts reads that overlap the interval ends (“overlap_interval_end_reads”, “span_interval_end_pairs”) or that map into an interval (“within_interval”) and calculates the median and mean coverage of the intervals. In case of junctions derived from alternative splice sites or exon skipping events the context sequence is divided into two intervals and overlap_interval_end reads represent the number of reads covering the junction of interest (“junction reads”). In case of intron retention, the interval is divided into three intervals. Here, the read coverage of the interval located in the middle represents the coverage of the intron of interest. **(B-C)** The distribution of **(B)** number of junction reads of junctions from A3SS, A5SS or ES events and **(C)** the median coverage of predicted retained introns.

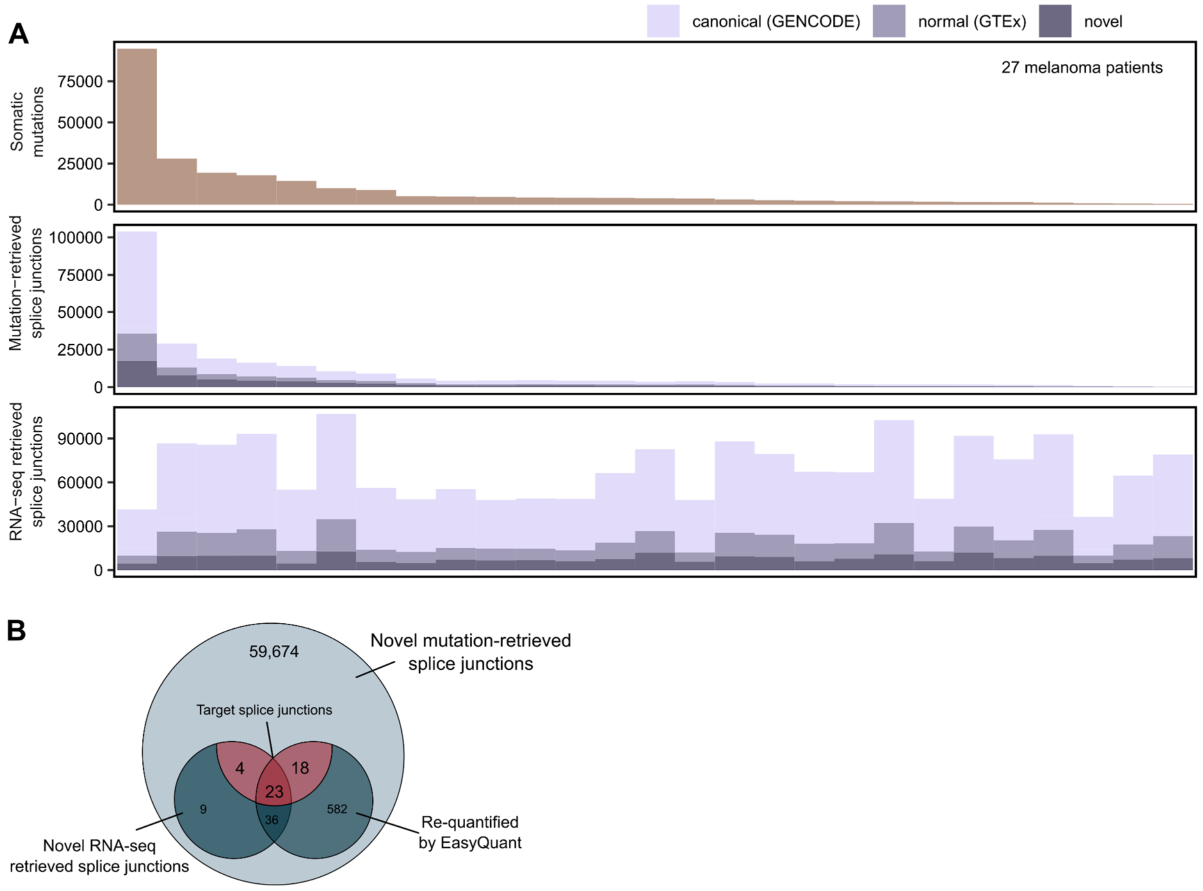


**Fig. S3: Overview of the verification set. (A)** The number of somatic mutations, mutation-derived splice junctions and RNA-seq derived splice junctions per sample in the verification cohort of 27 melanoma samples. **(B)** Overlap of mutation-retrieved splice junctions with those that were retrieved from RNA-seq by Spladder or Leafcutter, those that were re-quantified in RNA-seq with Easyquant and those that were predicted as targets.

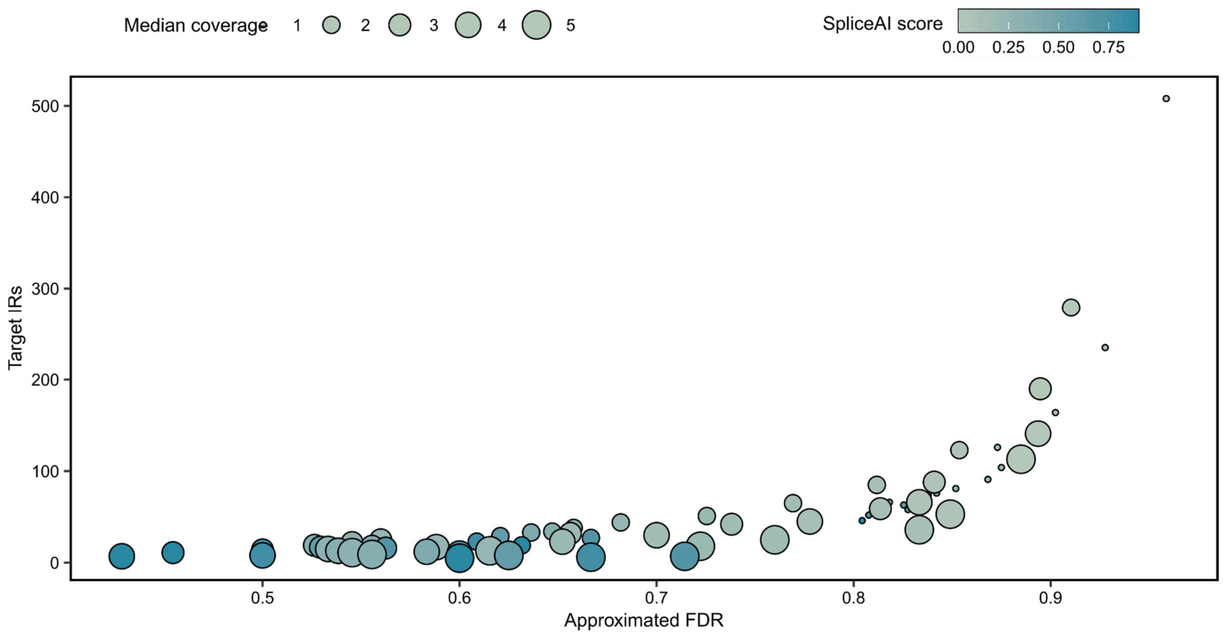


**Fig. S4: Estimation of false discovery rate (FDR) and number of targets from IR events.** Candidate IR events from the discovery were gradually filtered by thresholds on the mutation effect scores from and SpliceAI, and the re-quantification read support and the resulting approximated FDR and number of IR targets was determined.

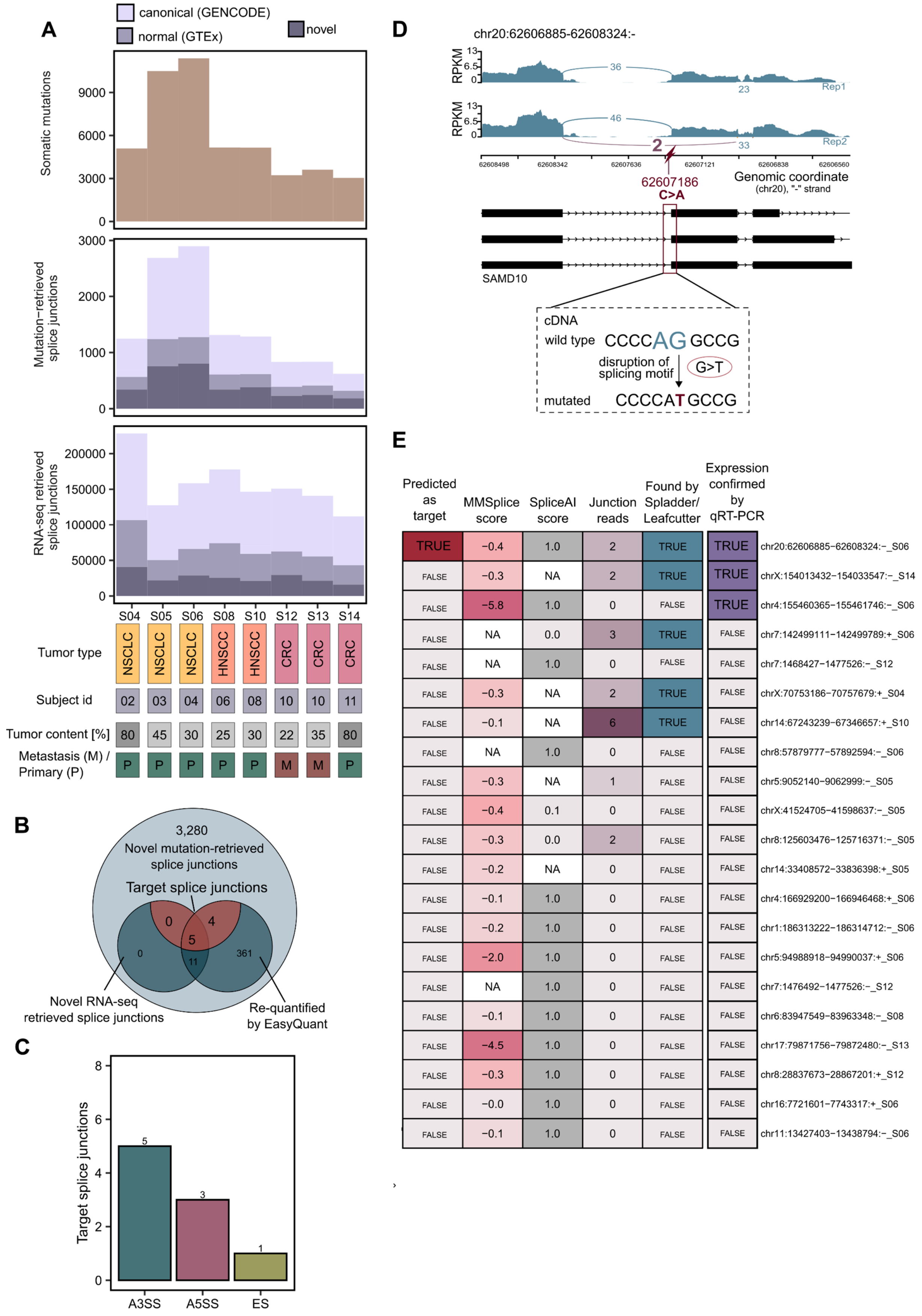


| Predicted as target | MMSplice score | SpliceAI score | Junction reads | Found by Spladder/ Leafcutter | Expression confirmed by qRT-PCR | |
|---|---|---|---|---|---|---|
| TRUE | −0.4 | 1.0 | 2 | TRUE | TRUE | chr20:62606885−62608324:−_S06 |
| FALSE | −0.3 | NA | 2 | TRUE | TRUE | chrX:154013432−154033547:−_S14 |
| FALSE | −5.8 | 1.0 | 0 | FALSE | TRUE | chr4:155460365−155461746:−_S06 |
| FALSE | NA | 0.0 | 3 | TRUE | FALSE | chr7:142499111−142499789:+_S06 |
| FALSE | NA | 1.0 | 0 | FALSE | FALSE | chr7:1468427−1477526:−_S12 |
| FALSE | −0.3 | NA | 2 | TRUE | FALSE | chrX:70753186−70757679:+_S04 |
| FALSE | −0.1 | NA | 6 | TRUE | FALSE | chr14:67243239−67346657:+_S10 |
| FALSE | NA | 1.0 | 0 | FALSE | FALSE | chr8:57879777−57892594:−_S06 |
| FALSE | −0.3 | NA | 1 | FALSE | FALSE | chr5:9052140−9062999:−_S05 |
| FALSE | −0.4 | 0.1 | 0 | FALSE | FALSE | chrX:41524705−41598637:−_S05 |
| FALSE | −0.3 | 0.0 | 2 | FALSE | FALSE | chr8:125603476−125716371:−_S05 |
| FALSE | −0.2 | NA | 0 | FALSE | FALSE | chr14:33408572−33836398:+_S05 |
| FALSE | −0.1 | 1.0 | 0 | FALSE | FALSE | chr4:166929200−166946468:+_S06 |
| FALSE | −0.2 | 1.0 | 0 | FALSE | FALSE | chr1:186313222−186314712:−_S06 |
| FALSE | −2.0 | 1.0 | 0 | FALSE | FALSE | chr5:94988918−94990037:+_S06 |
| FALSE | NA | 1.0 | 0 | FALSE | FALSE | chr7:1476492−1477526:−_S12 |
| FALSE | −0.1 | 1.0 | 0 | FALSE | FALSE | chr6:83947549−83963348:−_S08 |
| FALSE | −4.5 | 1.0 | 0 | FALSE | FALSE | chr17:79871756−79872480:−_S13 |
| FALSE | −0.3 | 1.0 | 0 | FALSE | FALSE | chr8:28837673−28867201:+_S12 |
| FALSE | −0.0 | 1.0 | 0 | FALSE | FALSE | chr16:7721601−7743317:+_S06 |
| FALSE | −0.1 | 1.0 | 0 | FALSE | FALSE | chr11:13427403−13438794:−_S06 |

**Fig. S5: Overview of the FFPE cohort. (A)** The number of somatic mutations, mutation-derived splice junctions and RNA-seq derived splice junctions per sample in the FFPE cohort. **(B)** Overlap of mutation-retrieved splice junctions with those that were retrieved from RNA-seq by Spladder or Leafcutter, those that were re-quantified in RNA-seq with Easyquant and those that were predicted as targets. **(C)** The number of target splice junctions from A3SS, A5SS and ES events. **(D)** Sashimi plot for the experimentally confirmed splice junction chr20:62606885-62608324:- from an ES event that was predicted as a target. The target splice junctions may be caused by a SNV that changes C to A on the negative strand, resulting in a disruption of an acceptor motif in the cDNA. **(E)** The RNA expression of 21 splice junctions from ES events were analyzed with qRT-PCR. Heatmap showing MMSplice and SpliceAI score, the number of junction reads, whether a junction was found by an RNA-seq tool (Leafcutter/Spladder) and whether junctions were predicted as targets.
